# pGG-PIP: A GreenGate (GG) entry vector collection with Plant Immune system Promoters (PIP)

**DOI:** 10.1101/2022.12.20.521163

**Authors:** Jacob Calabria, Madlen I. Rast-Somssich, Liu Wang, Hsiang-Wen Chen, Michelle Watt, Alexander Idnurm, Staffan Persson, Marc Somssich

## Abstract

The regulatory sequences controlling the expression of a gene (i.e., the promoter) are essential to properly understand a gene’s function. From their use in mutant complementation assays, to studying their responsiveness to different stimuli via transcriptional reporter lines or using them as proxy for the activation of certain pathways, assays using promoter sequences are valuable tools for insight into the genetic architecture underlying plant life. The GreenGate (GG) system is a plant-specific variant of the Golden Gate assembly method, a modular cloning system that allows the hierarchical assembly of individual donor DNA fragments into one expression clone via a single reaction step. Here, we present a collection of 75 GG entry vectors carrying putative regulatory sequences for *Arabidopsis thaliana* genes involved in many different pathways of the plant immune system, designated Plant Immune system Promoters (PIP). This pGG-PIP entry vector set enables the rapid assembly of expression vectors to be used for transcriptional reporters of plant immune system components, mutant complementation assays when coupled with coding sequences, mis-expression experiments for genes of interest, or the targeted use of CRISPR/Cas9 genome editing. We used pGG-PIP vectors to create fluorescent transcriptional reporters in *A*. *thaliana* and demonstrated the potential of these reporters to image the responsiveness of specific plant immunity genes to infection and colonization by the fungal pathogen *Fusarium oxysporum*. Using the PLANT ELICITOR PEPTIDE (PEP) pathway as an example, we show that several components of this pathway are locally activated in response to colonization by the fungus.

## Introduction

Since the development of molecular cloning in the early 1970s, the isolation of genes and promoters, and subsequent transgenesis of model organisms, has become standard practice in the life sciences (Somssich, 2022). The development of more advanced cloning methods, particularly the recombination-based Gateway technology in the year 2000, made the creation of expression clones ready for transformation ever easier (Hartley *et al*., 2000). However, with the rise of the synthetic biology field, it is now no longer sufficient for these methods to facilitate the cloning of individual DNA fragments. To recreate entire pathways and gene circuits in plants and other model organisms, larger DNA constructs need to be readily assembled from individual components, and these distinct building blocks need to be compatible to allow the flexibility to mix and match different promoters, coding sequences, protein tags, terminators, and resistance genes for selection of transgenic lines (Meng and Ellis, 2020). This requirement was met with the new modular cloning techniques which use recombination-based hierarchical assembly of multiple donor- modules (each containing, for example, a promoter of choice, tag of choice, gene of interest, etc.) into one ordered expression clone to be used for transgenesis (Fig. 1) (Bird *et al*., 2022). Among the developed modular cloning methods, the Golden Gate system has emerged as the most widely utilized version, and in 2013 Lampropoulos *et al*. developed the plant-specific GreenGate variant of the Golden Gate technique (Engler *et al*., 2008, 2009; Weber *et al*., 2011; Lampropoulos *et al*., 2013). The GreenGate toolkit provides users with a wide range of entry vectors that serve as donors for standard promoters (e.g. CaMV35S, UBQ10), protein tags (e.g. GFP, NLS, HDEL), terminators (e.g. CaMV35S, UBQ10) and plant resistance cassettes (e.g. BastaR, HygR, KanR), to build basic gene expression modules for plants.

**Figure 1:**
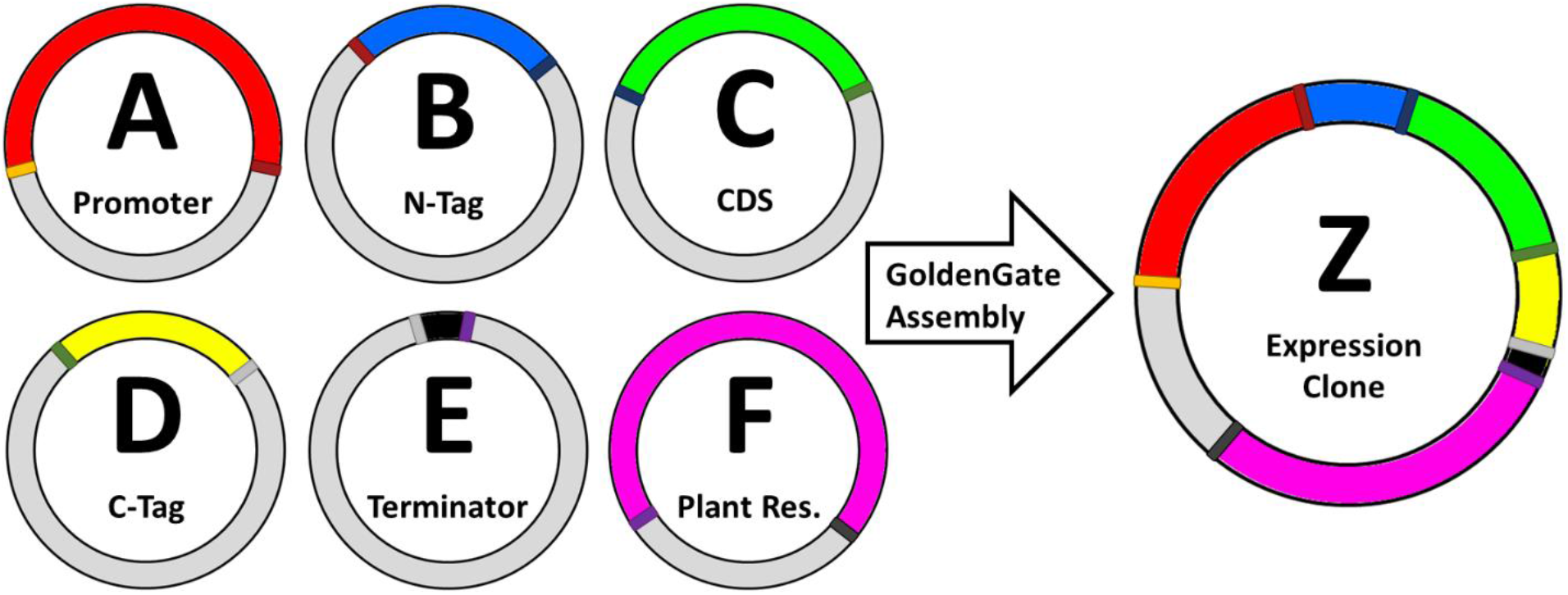
Overview of the modular Golden/GreenGate cloning principle. The promoter, N- and C-terminal tags, the gene-of-interest coding sequence (CDS), a terminator, and a plant resistance cassette are all cloned into individual entry clones (A-F), which are then combined with an ‘empty’ destination clone (Z) in the Golden Gate assembly mix. During the assembly reaction, the *Bsa*I restriction enzyme cuts the different fragments from their respective entry clones. These fragments subsequently self-align via complementary four base pair overhangs (indicated by the thin colored borders on each entry vector; complementary overhangs that will align have the same color), and the T4 DNA-ligase fuses the individual fragments in the final expression clone (Z) that can be used to transform plants.

Additional kits have been developed and added to the GreenGate toolbox since 2013, providing entry vector sets to build plasmids for CRISPR/Cas9-guided genome editing (Wu *et al*., 2018), CRISPR/Cas9-guided tissue-specific gene knockout (Decaestecker *et al*., 2019) and inducible and cell-type specific gene expression (Schürholz *et al*., 2018). In addition, a toolbox of fluorescent proteins suitable for work in plants (Denay *et al*., 2019), along with the necessary clones for *in planta* proximity ligation assays using the biotin ligase (Goslin *et al*., 2021), and destination vectors with an already integrated plasma membrane-marker (Kümpers *et al*., 2022) have been designed. The development and availability of these various toolboxes, all compatible with each other, demonstrate the usefulness of such modular systems to enable researchers to quickly adopt new technologies, providing the flexibility and versatility to combine and recombine their existing vectors (i.e., modules) with new vectors from all other kits.

In our work we use a microscopy-based live-imaging approach to monitor the *Arabidopsis thaliana*’s defense responses to infection and colonization by the pathogenic fungus *Fusarium oxysporum* strain *Fo*5176 (*Fo*5176) on an individual cell level (Calabria *et al*., 2022; Wang *et al*., 2022*a*). Fluorescent transcriptional reporters are a good tool for such studies, as activation of certain pathways is typically associated with transcriptional upregulation (Ngou *et al*., 2021). However, most studies investigating the transcriptional responsiveness of certain pathways to pathogens tend to use transcriptomic analyses of whole tissues, organs, or seedlings, therefore losing all spatial resolution and differences between cell groups and even individual cells. The responses of only the few cells that could be the major contributors to a host-microbe outcome are also typically lost in such experiments, as they are below the threshold of background noise. A microscopy-based approach that allows for the spatial resolution and observation of responses in individual cells therefore closes this knowledge gap. One reason why such an approach is not more common is that a prerequisite for this work is the availability of fluorescent reporter lines for all the different plant immune pathways to be investigated. To this end, we have cloned the putative regulatory sequences (i.e., promoters) of 75 *A. thaliana* genes, representing many of the major branches of the plant immune system (Fig. 2 and Table 1). We have done so using the GreenGate cloning system, therefore creating a GreenGate plant promoter entry vector set that is compatible with all other GreenGate-based toolkits, and that we believe will be a valuable tool for the scientific community.

**Figure 2:**
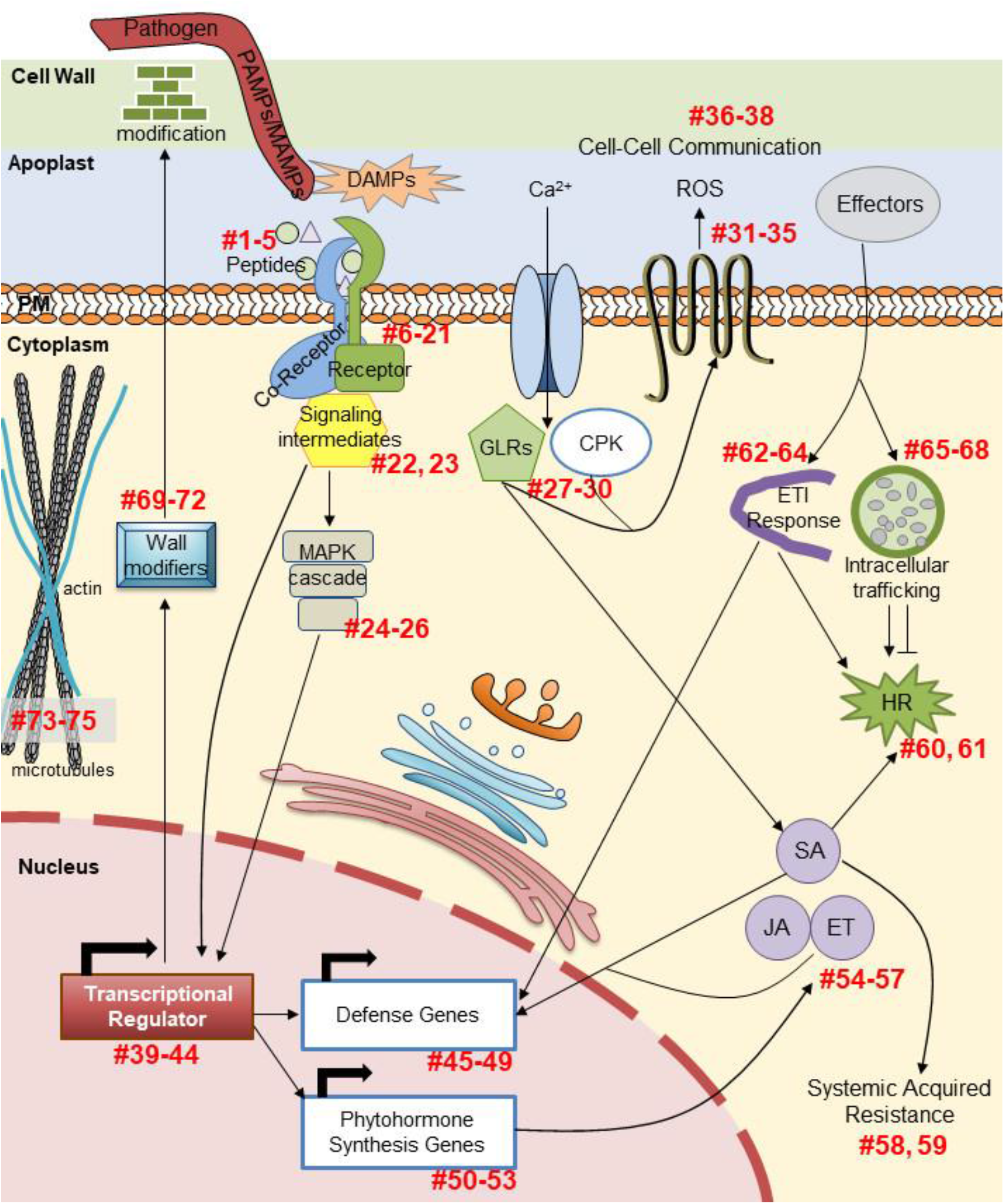
Overview of the different immune pathways represented in the pGG-PIP collection. The red numbers correspond to the pGG-PIP plasmid numbers in column one of Table 1.

**Table 1:**
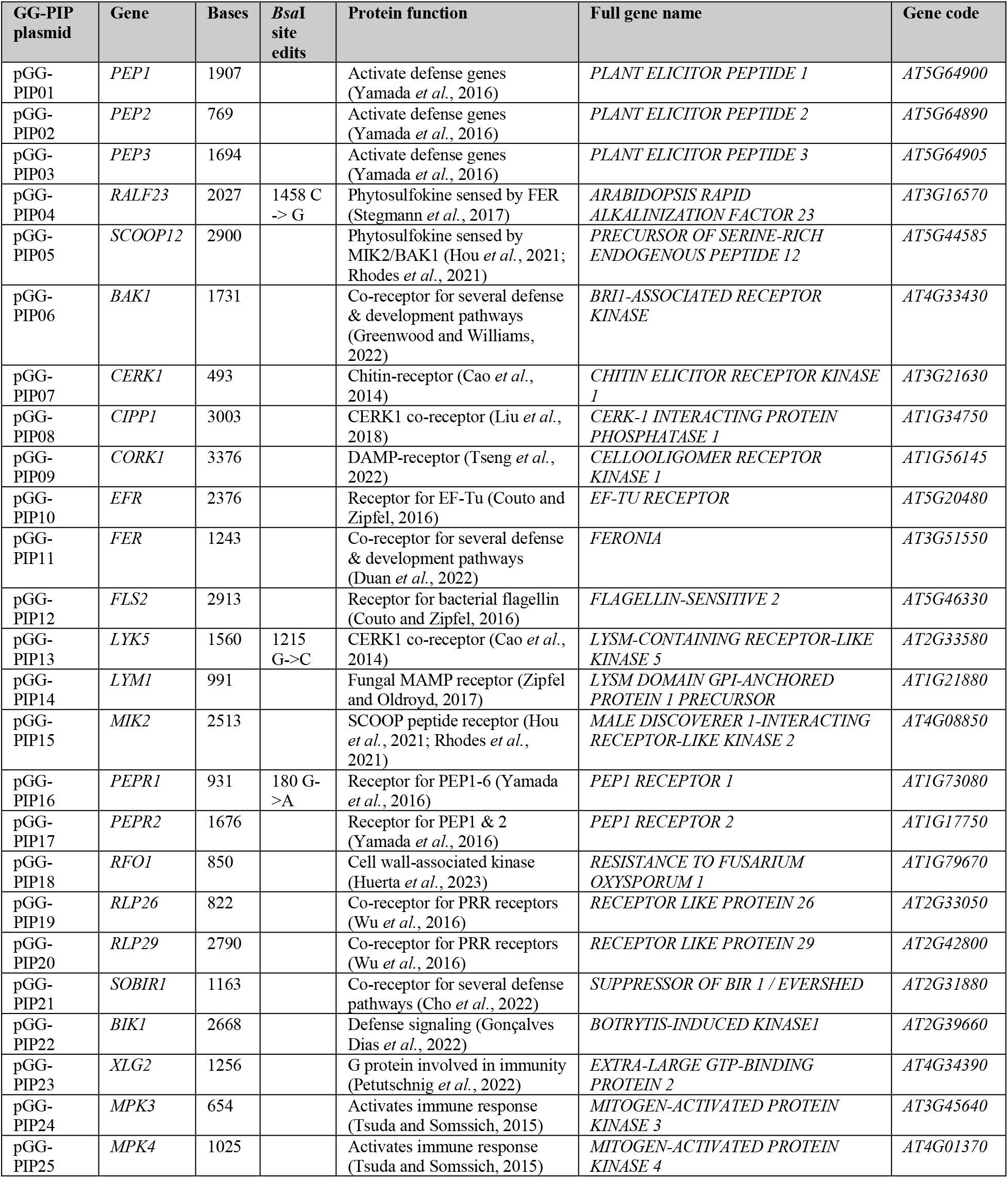
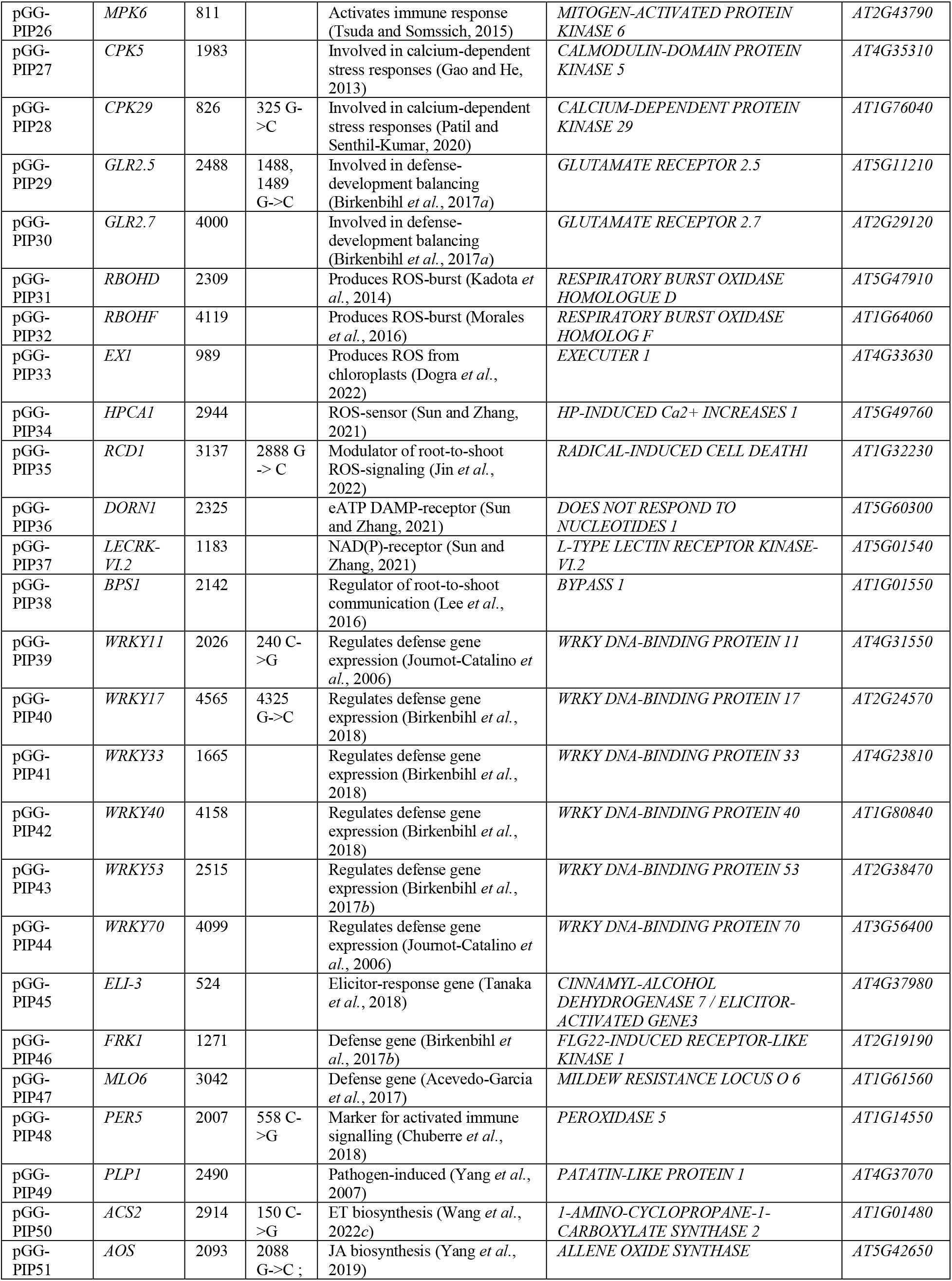
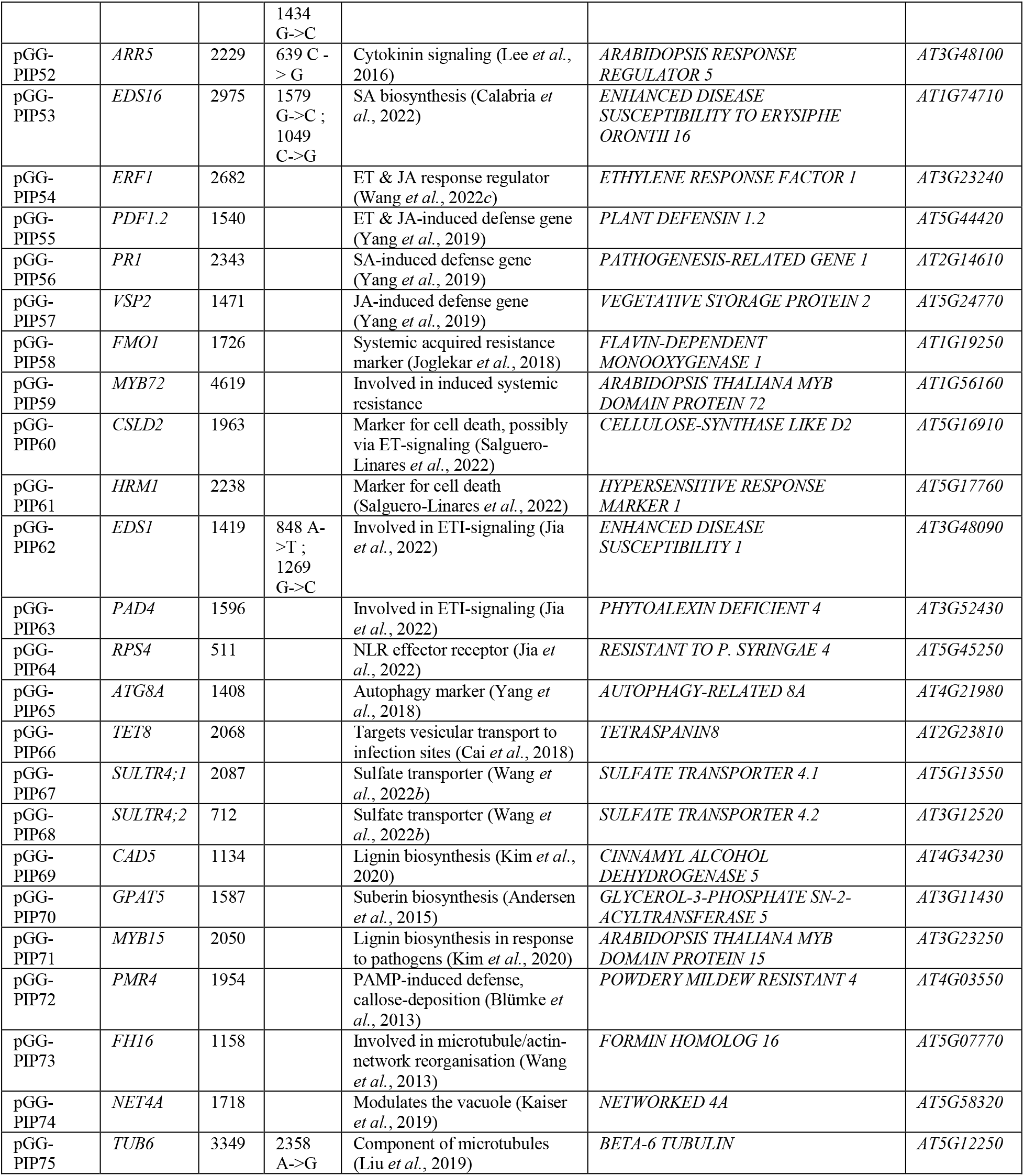
List of the 75 pGG-PIP promoter entry vectors in the set described in this paper. The pGG-PIP numbers correspond to the red numbers in Fig. 2.

As a proof-of-principle, we selected a number of these promoters to create fluorescent transcriptional reporter lines for genes involved in the PLANT ELICITOR PEPTIDE (PEP)- pathway. The PEP pathway is involved in the immune response to several plant pathogens, including bacteria, fungi and oomycetes (Saijo *et al*., 2018). The PEP1-6 peptides function as phytosulfokines. They are expressed as part of the plant’s pattern-triggered immunity (PTI) response and trigger cell autonomous and non-autonomous defense responses when they are perceived by the plasma membrane localized receptors PEPR1 and PEPR2 (Yamaguchi *et al*., 2006, 2010; Saijo *et al*., 2018). For their function, they require the intracellular kinase BIK1, which activates the NADPH oxidase RBOHD to trigger a ROS-burst, and also induces defense gene expression (Liu *et al*., 2013; Kadota *et al*., 2014; Saijo *et al*., 2018; Jing *et al*., 2020). A role for the PEP pathway in the defense against fungal disease has been hypothesized, based on large-scale transcriptomic data. Plants carrying mutations in *MEDIATOR18* and *20* show increased resistance to infection by *F. oxysporum* strain *Fo*5176, and mRNA sequencing has demonstrated that *PEP1* is among the genes no longer induced after infection of these mutant plants (Fallath *et al*., 2017; Wang *et al*., 2022*a*). Similarly, *PEPR1* and *2* were among the genes suppressed by the endophytic *F*. *oxysporum* strain *Fo*47, indicating that the PEP pathway normally acts to restrict fungal colonization (Guo *et al*., 2021; Wang *et al*., 2022*a*). However, no detailed data is available. Here, we show that colonization of the *A. thaliana* vasculature is accompanied by a highly localized activation of this pathway via upregulation of genes encoding the peptide ligand PEP1, as well as the cognate PEP RECEPTORs 1 and 2, the downstream signaling intermediate BIK1 and the NADPH oxidase RBOHD.

## Results

### Selection and cloning of the promoters

By using these putative promoter sequences for the construction of transcriptional reporter lines, our main aim is to obtain information about the specific localized activation of different immune pathways in response to infection by a pathogen, in our case *Fo*5176. Once we have identified pathways involved in the plant’s defense against this strain, we plan to further extend these experiments based on what is known about the function of these pathways. Thus, it is important to select promoters from many different pathways of the plant immune system, but also from pathways that are particularly well understood, since this existing knowledge will guide our future work. Some of the best studied immune pathways in the plant are the pattern-recognition flagellin (flg) and elongation factor thermos unstable (EF-Tu) pathways, the phytohormones jasmonic acid (JA), salicylic acid (SA) and ethylene (ET), and the transcriptional activity of several WRKY transcription factors. We therefore selected representative gene promoters from such well understood pathways, but also additional candidates such as MAP and calcium-dependent protein kinases (MPKs, CPKs), genes involved in production and signaling via reactive oxygen species (ROS), and genes involved in effector-triggered immunity (ETI) responses via the EDS1-PAD4 module. Figure 2 and Table 1 provide a complete overview of the 75 promoters included in the pGG-PIP collection. The numbers in Fig. 2 correspond to the pGG-PIP plasmid number in the first column of Table 1.

For the construction of the pGG-PIP plasmids, we considered the entire stretch of DNA from the 3° upstream neighboring gene’s coding sequence to the ATG of the gene of interest as the gene’s putative regulatory sequences, and thus its ‘promoter’. There are a few exceptions to this rule, where we used a defined stretch of DNA that has previously been described to complement a mutant (e.g., *FMO1*, *RPS4* (Wirthmueller *et al*., 2007; Joglekar *et al*., 2018)). Where present, this may include untranslated regions (UTRs), pseudogenes, or transposons, since these elements could indeed affect expression of a gene *in planta*. For the recombination-based cloning, the *Bsa*I recognition site and standard four base pair GreenGate A overhangs (5°-ACCT and TTGT-3°) were added via primers during amplification of the promoter sequences (see Supplementary Table 1 for a list of primer sequences). If the promoter sequence contained an internal *Bsa*I restriction site (GGTCTC), we mutated a single base, usually a G to a C via scar-free *Bsa*I cloning, i.e., we inserted the base mutation by making it part of the four base pair overhang used for the scar-free cloning (see Table 1 for the exact sequence edits made, and Supplementary Table 1 for primer sequences). The promoters were then transferred into the GreenGate promoter entry vector pGGA000 via the four base pair A overhangs in a Golden Gate assembly reaction.

### Using pGG-PIP vectors to create transcriptional reporters for the PEP pathway

To test the usefulness of these promoters to map local individual cell responses upon attack by a microbe, we created transcriptional reporter lines for genes involved in the PEP pathway. We used a nuclear localized tandem of mTurqoise2 (mT2) fluorophores as a reporter, since mT2 has been shown to be very bright and photostable in plant cells (Goedhart *et al*., 2012; Long *et al*., 2018; Denay *et al*., 2019). The nuclear localization enhances the brightness even further due to molecular crowding, and furthermore helps to identify individual cells.

We first probed our reporters for the PEP1 and PEP2 peptides (pGG-PIP01 and 02), as well as the PEPR1 and PEPR2 receptors (pGG-PIP16 and 17), assessing their local responsiveness to colonization by *Fo*5176. Under control uninfected conditions, *PEP1* was weakly expressed in inner tissue cells of the root differentiation zone (DZ) (Fig. S1A-D). No expression was detectable in the root tip, the meristematic or elongation zones (MZ and EZ). *PEP2* was robustly expressed in the DZ, but in contrast to *PEP1*, its expression was stronger in the outer tissues (Fig. S1G-H). In addition, it was also expressed in the root tip, around the MZ and early EZ (Fig. S1E-F). *PEPR1* was generally expressed at a very weak level. In the root tip, starting with the EZ, expression was limited to the vasculature, while from the young DZ (or root hair zone) onward, expression was restricted to the outer tissues (Fig. S2A-D). *PEPR2* was expressed only in the DZ, and only in the vasculature, but expression was much stronger compared to *PEPR1* (Fig. S2E-H). Thus, in vascular cells, *PEPR1* and *PEPR2* complemented each other under uninfected conditions. Following colonization of the vasculature by *Fo*5176, *PEP1*, as well as both *PEPR*s, showed a strong upregulation in the cells adjacent to the colonization site (Fig. 3). Interestingly, this upregulation was limited to the cells of the vasculature, the exact tissue that is targeted and colonized by *Fo*5176, showing how precisely the plant targets its response. Importantly, the induction of *PEPR2* was much stronger than the induction of *PEP1* and *PEPR1* (Fig. 3G-J). Interestingly, *PEP2* also showed an induction next to the colonized tissue, but its expression appeared to be limited to the outer tissues, the epidermis and cortex, rather than the colonized vasculature (Fig. 3C, D). The strong induction of the *PEPs* and *PEPRs* remained visible for roughly 10 to 12 cells, and then tapered out, eventually returning to the expression pattern of the uninfected control.

**Figure 3:**
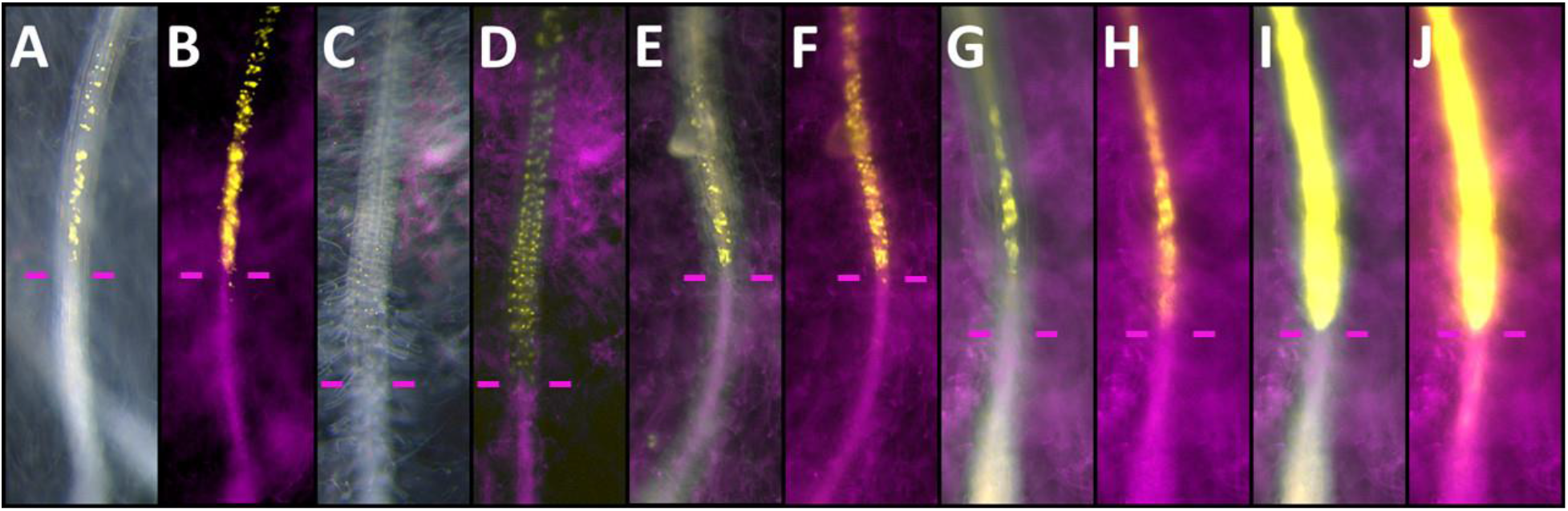
Local transcriptional responses in *PEP1*, *PEP2*, *PEPR1* and *PEPR2* expression to colonization by *Fo*5176. Bright field and fluorescence (A, C, E, G, I) and fluorescence only (B, D, F, H, J) images of colonized roots expressing the *PEP1* (A, B), *PEP2* (C, D), *PEPR1* (E, F) or *PEPR2* (G-J) reporters. *Fo*5176 is shown in magenta, the reporters in yellow. The purple bars indicate the fungal colonization front. All images were recorded with comparable imaging settings, except for G and H, which are the same as I and J, just with reduced exposure time. Expression of the *PEPR2* reporter was so strong (I, J), that we reduced the exposure time to visualize individual cells in G and H.

Since it was shown that the ability of the PEPs and PEPRs to induce any signaling downstream of the receptors is strictly dependent on the activity of the cytoplasmic kinase BIK1, we also investigated the responsiveness of a *BIK1* (pGG-PIP22) transcriptional reporter (Liu *et al*., 2013; Yamada *et al*., 2016). Under uninfected control conditions, *BIK1* is expressed in the MZ and DZ, with stronger expression in the outer tissues, and only weak expression in the vascular tissue of the MZ, somewhat resembling the expression pattern of *PEP2* (compare Fig. S3A-D and S1E-H). Following colonization of the root by *Fo*5176, this pattern changed significantly, with *BIK1* now expressed specifically in the cells immediately bordering the colonized tissue, with a clearly visible expression maximum in the vasculature (Fig. 4A, B). Hence, the expression maxima of *BIK1*, *PEP1*, *PEPR1* and *PEPR2* all overlapped in a small group of vascular cells closest to the colonized tissue. *BIK1* however, also showed expression in the endodermis and cortex, albeit at a much lower level than in the vasculature, while *PEP1* and *PEPRs* expression were restricted to the vasculature (Fig. 3 and Fig. 4A, B).

**Figure 4:**
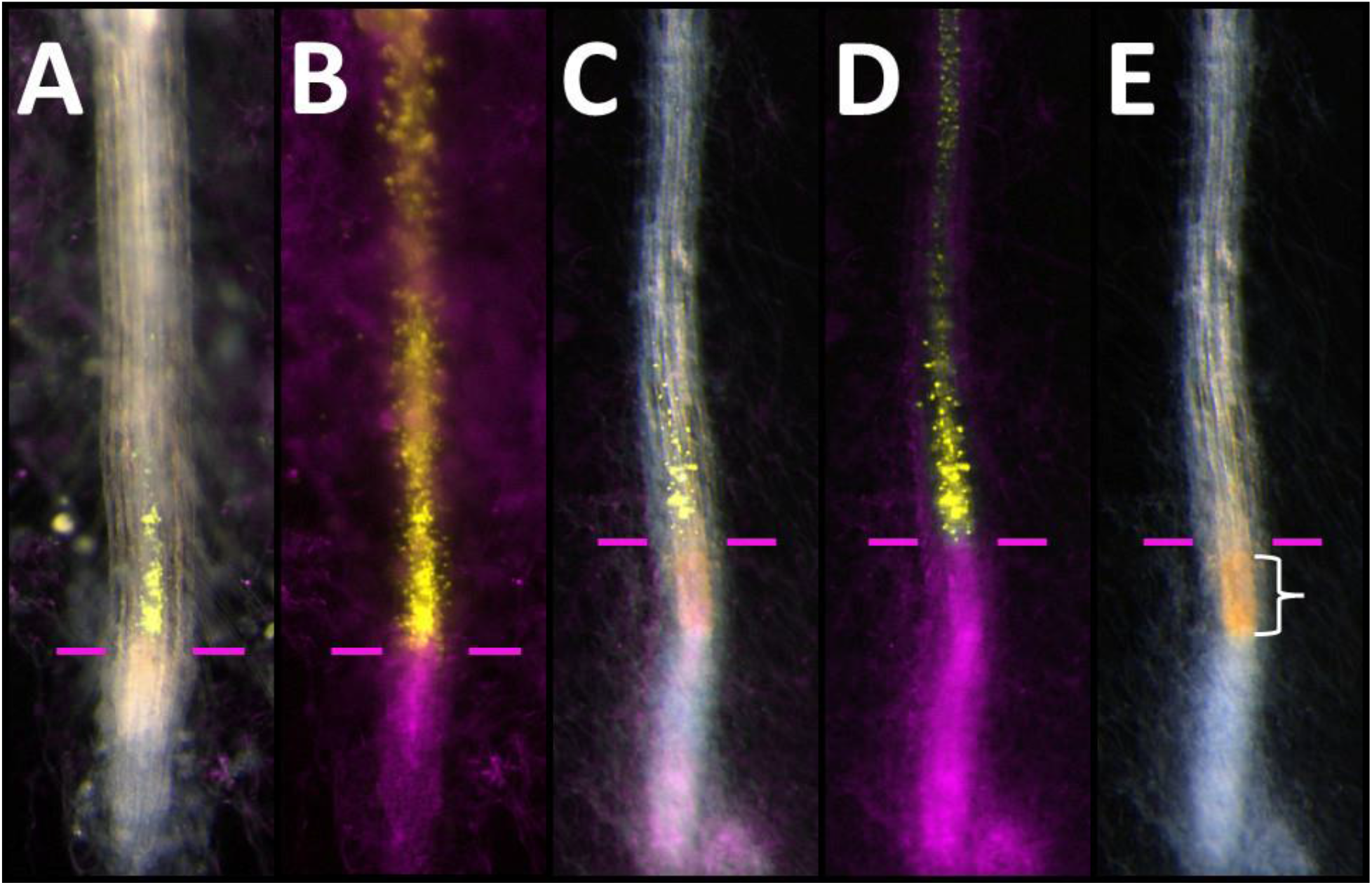
Transcriptional responses of *BIK1* and *RBOHD* expression to colonization by *Fo*5176. Bright field and fluorescence (**A, C**), fluorescence only (**B, D**), or bright field only (**E**) images of a colonized root expressing the *BIK1* (A, B), or *RBOHD* (C, D, E) reporters. The *Fo*5176 is shown in magenta, the reporters in yellow. The purple bars indicate the fungal colonization front. The white bracket in E indicates the area with ‘root browning’.

One of the downstream outputs of PEP-activated BIK1 is the activation of the NADPH oxidase RBOHD to trigger an apoplastic ROS-burst (Holmes *et al*., 2018; Jing *et al*., 2020). We therefore investigated if *RBOHD* (pGG-PIP31) is also activated in the cells expressing *PEP1*, the *PEPRs* and *BIK1*. Under uninfected control conditions, *RBOHD* is expressed in all cells and tissues of the DZ, but not in the root tip (MZ, EZ) (Fig. S3E-H). Upon infection by *Fo*5176, expression is activated in the cells next to the colonization site, even if they are still part of the EZ, which normally does not express *RBOHD* (Fig. 4C, D). The pattern we observed very much resembled the pattern of *BIK1* expression in response to colonization, with a maximum in a small group of vascular cells next to the colonization site (Fig. 4). However, in contrast to the expression of *BIK1* and the *PEPs* and *PEPRs*, upregulation of *RBOHD* appeared to be more spatially restricted to the cells closest to the colonization site. *RBOHD* expression quickly returned to control levels behind this small group of cells, while the activation of the other markers appeared to taper out more gradually, across a longer stretch of cells.

While imaging the fungal infected reporter lines, we also regularly observed a discoloration of the root in the colonized tissue (white bracket in Fig. 4E). This has previously been described as ‘root browning’, and has been observed in response to infection by *F. oxysporum* or treatment of the plant with SERINE-RICH ENDOGENOUS PEPTIDEs (Tintor *et al*., 2020; Hou *et al*., 2021). While it remains unclear what the exact cause for this discoloration is, redox/oxidative stress induced by ROS is one possibility. In our assay, the browning does not completely overlap with the expression maximum of *RBOHD*, but as the colonization of the fungus progresses upward through the vasculature, the *RBOHD* expression also extends along adjacent to the colonization front. Therefore, the area showing root browning would be the region that last expressed *RBOHD* at high levels, and thus could have been under redox/oxidative stress, but this requires further examination.

## Discussion

In this paper we present a GreenGate-based entry vector set with 75 plant immune system promoters. In combination with the basic GreenGate kit from Lampropoulos et al. (2013) this is already sufficient to build simple tools, such as transcriptional reporter lines, that can then be probed for their responsiveness to a pathogen, elicitor, or other stimulant. Furthermore, owing to the universal compatibility of these entry vectors with all other GreenGate-based systems, this vector set can be used to screen for interactors of proteins in their native expression domain using the proximity ligation kit from Goslin *et al*. (2021), or for tissue-specific gene knockouts by combining them with the CRISPR-TSKO kit from Decaestecker *et al*. (2019), to name just two possibilities.

As a straightforward proof-of-principle, we used some of the pGG-PIPs to create fluorescent reporter lines for the PEP pathway and demonstrate how these can be used to gain insights into *spatial immunity* – the highly localized activation of an immune pathway. Using these lines, we showed that *PEP1*, and to some degree *PEP2*, were responsive to infection and colonization by the vascular pathogen *Fo*5176. The danger signal PEP1 was upregulated in a specific group of vascular cells immediately adjacent to the fungal colonization site, as were the two receptors *PEPR1* and *2*. We thus hypothesize that activation of the PEP pathway is part of the plant’s immune response to infection and colonization by *Fo*5176. Importantly, we could show the highly targeted activation of this pathway in just a couple of vascular cells immediately adjacent to a colonization site. Such a local response has rarely been shown *in vivo* and *in planta* before, and is typically overlooked in large-scale transcriptomic analyses.

Since the PEPRs are unlikely to serve as pattern-recognition receptors (PRRs) to sense the pathogen, it is more likely that the PEPRs are part of a larger complex containing the PRR co-receptor BAK1, as well as other PRRs, such as MIK2 or FER, since the interaction between BAK1 and the PEPRs has been previously shown, and all of these PRRs have been implicated to function in the defense against *Fo*5176 (Yamada *et al*., 2016; Wang *et al*., 2022*a*) (Fig. 5). Possibly, these co-receptors are all co-localized in plasma membrane nanodomains to facilitate the rapid assembly of larger receptor platforms (Somssich *et al*., 2015; McKenna *et al*., 2019; Somssich, 2020; Gronnier *et al*., 2022). Indeed, rather than functioning as a PRR sensing the pathogen directly, work from a recent publication suggests that the PEPRs act as extracellular pH sensors (Liu *et al*., 2022). Alkalinization of the apoplast is a hallmark of activated immunity, and Liu *et al*., (2022) show that the PEPRs specifically interact with their co-receptor BAK1 under alkaline conditions, while they only show a weak affinity for interaction under acidic conditions (Felix *et al*., 1993; Liu *et al*., 2022). Interestingly, *F. oxysporum* uses functional homologs of plant RAPID ALKALINIZATION FACTORs to induce alkalinization of the apoplast, as this also stimulates infectious growth of the fungus (Masachis *et al*., 2016; Wang *et al*., 2022*a*). Thus, the PEPRs pathway may act as pH sensors to counteract this action by the fungus, improving the plant’s defense under these alkaline conditions. Extracellular binding of the PEPs to their receptors generally leads to the activation of the PEPRs intracellular kinase domain and trans- phosphorylation of the interacting cytoplasmic kinase BIK1 (Liu *et al*., 2013). Indeed, we could show that our *BIK1* reporter is also activated in the same pattern as the *PEP1* and the *PEPR* reporters. BIK1 activation then facilitates downstream intracellular signaling (Yamada *et al*., 2016). These downstream signaling events eventually result in the transcriptional activation of defense genes, but also auto-activation of the *PEP* genes (Fig. 5). Since the PEPs do not have an identifiable secretion signal that would direct them into the secretory pathway, it is assumed that they are released into the apoplast indirectly when a pathogen breaches the plasma membrane and causes local membrane disintegration (Fig. 5) (Yamaguchi and Huffaker, 2011; Yamada *et al*., 2016). Once in the apoplast, they are then perceived by the PEPRs, creating a positive feedback-loop of PEP-signaling and an auto-amplification of the plant’s defense response (Fig. 5). A second intracellular pathway activated via BIK1 leads to activation of the NADPH oxidase RBOHD (Fig. 5) (Kadota *et al*., 2014; Holmes *et al*., 2018; Jing *et al*., 2020). This pathway was proposed to be primarily driven by PEP1 and the PEPR2 (Jing *et al*., 2020) and indeed, *PEP1* and *PEPR2* are also the two reporters showing the strongest response to colonization by *Fo*5176, while *PEP2* was primarily upregulated in the cells around the vasculature. *PEPR1* was also activated in the same pattern as *PEP1*, *PEPR2*, and *BIK1*, but at a much lower level compared to *PEPR2*. Further, we could also confirm that *RBOHD* expression is activated in these same cells, indicating that this *BIK1*-*RBOHD*-dependent pathway for ROS-release is also potentially involved in the defense against *Fo*5176 (Fig. 5). The activation of RBOHD, and the resulting ROS-burst, would aid in the plant’s defense against *Fo*5176, with ROS being directly harmful to the pathogen. Additionally, ROS is known to also function as a signal in cell-cell communication to prime the surrounding tissue for impeding attack (Waszczak *et al*., 2018). We believe that the local ‘root browning’ that we observed in the infected roots is likely the result of this apoplastic ROS-burst.

**Figure 5:**
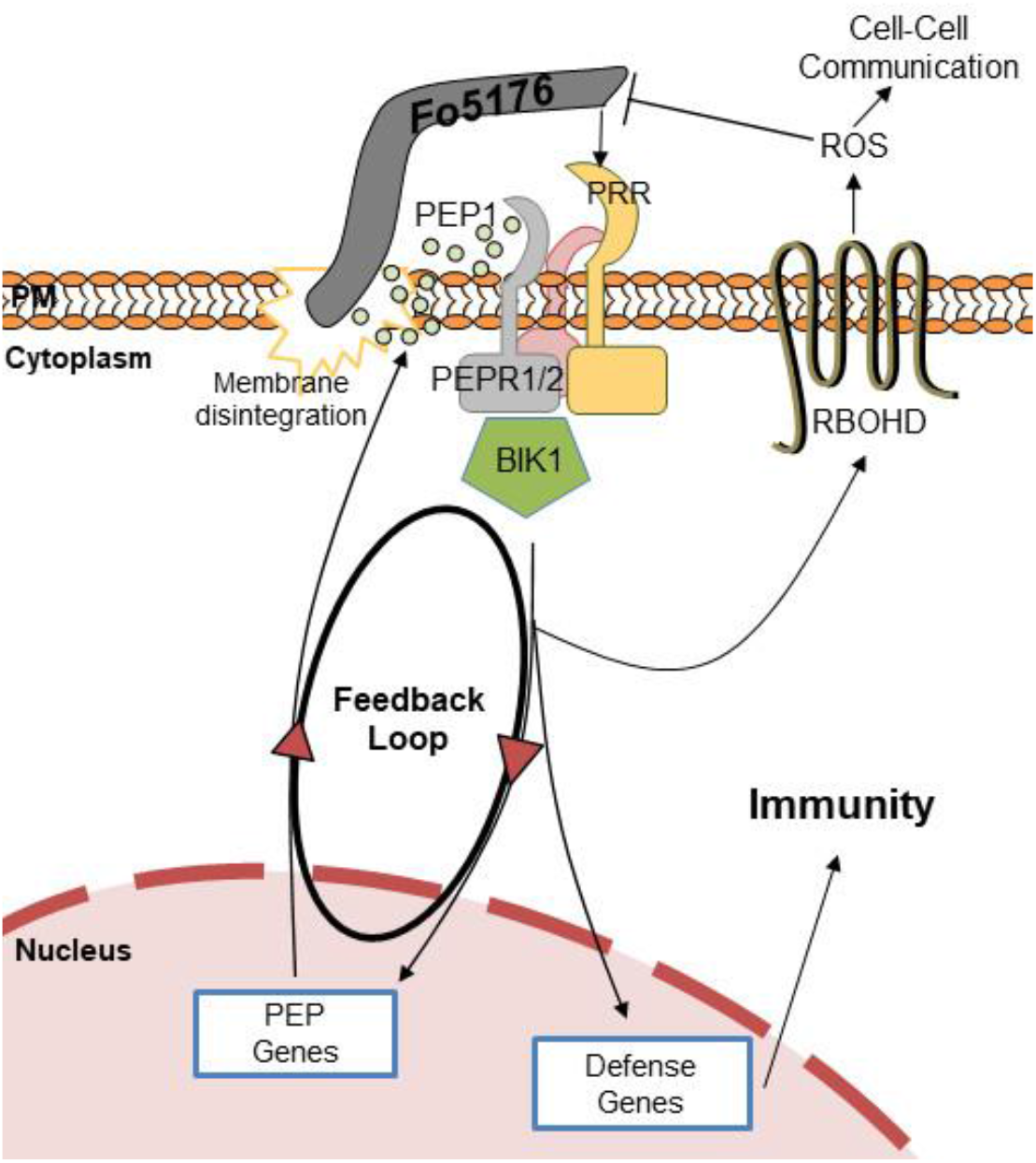
Model of the PEP pathway in response to *Fo*5176. Alkalinization of the apoplast via *Fo*5176- derived alkalinization factors induces interaction of the PEPRs with their PRR co- receptors. Intracellularly the receptors phosphorylate and activate BIK1. BIK1 activates expression of PEP1, thereby amplifying the PEP signal and creating a positive feedback loop. BIK1 also activates other defense genes, possibly via a MAPK cascade, and activates RBOHD to produce an apoplastic ROS-burst, which is damaging to the pathogen, and functions as cell-cell communication signal.

A role for the PEP-pathway in the defense against *F. oxysporum* has previously been hypothesized on the basis of transcriptomic data showing up- or downregulation of some of the components in response to infection (Fallath *et al*., 2017; Guo *et al*., 2021; Wang *et al*., 2022*a*). By investigating the responsiveness of our transcriptional reporters for PEP1 and 2, as well as PEPR1 and 2, we added spatial resolution to the available transcriptomic data. We could show that PEP1 and the PEPRs act in a clearly defined group of vascular cells immediately next to the fungal colonization site, and this localized response overlaps with the activation of downstream signaling factors and a local ROS-burst. *PEP2* was also induced in response to colonization, but it appeared to be upregulated in the outer tissues, not the infected vasculature. While this may indicate a role for PEP2 in priming these neighboring tissues, the fact that neither of the two *PEPRs* were expressed there, makes it unclear how PEP2 would be perceived in these cells. Importantly, when Fallath *et al*. (2017) investigated the transcriptional response of *med* mutants to infection by *Fo*5176, they also only noted a deregulation of *PEP1* in these mutants, but not *PEP2*, indicating that *PEP1* is indeed the major PEP signal acting in response to colonization by *Fo*5176.

Finally, we believe that the pGG-PIP plant immune promoter resource introduced in this article will be valuable and helpful to the community for various research approaches. We will make the 75-plasmid collection available via AddGene for quick and simple distribution to the community. We aim to provide further detailed ‘spatial immunity’ information by using these pGG-PIPs in our future work.

## Material & Methods

### Cloning of the pGG-PIPs and transcriptional reporter constructs

The pGG-PIP entry vectors are based on the pGGA000 vector from the original GreenGate kit (Lampropoulos *et al*., 2013). The different promoters were amplified from total cellular *A. thaliana* (natural accession Columbia) DNA, extracted from rosette leaves with the Qiagen DNeasy Plant Kit, and using the primers in supplementary table 1. For error-free amplification, the Phusion high-fidelity DNA polymerase with proofreading (New England Biolabs) was used. The fragments were then transferred into pGGA000 in a Golden Gate assembly reaction using the NEBridge Golden Gate Assembly Kit (BsaI-HF v2) (New England Biolabs). To clone the transcriptional reporter lines, we used the destination vector pGGZ003, as well as the donor vectors pGGD007 (*linker-NLS* (Nuclear Localization Signal)), pGGE009 (*UBQ10* terminator), and pGGF005 (*pUBQ10::HygR:tOCS*) from the original GreenGate kit from Lampropoulos *et al*. (2013). pGGB- mT2 (*mTurquoise2*) and pGGC-mT2 (*mTurquoise2*) were created by cloning the *mT2* coding sequence from an AddGene-derived template into pGGB000 and pGGC000 from the original GreenGate kit (Goedhart *et al*., 2012; Lampropoulos *et al*., 2013). In combination with our pGG- PIPs, this yielded the *pPIP::mT2-mT2-NLS:tUBQ10:pUBQ10::HygR:tOCS* reporters.

### Plant growth and transformation

*A. thaliana* Columbia plants (Somssich, 2018) were grown at 16-hour light conditions, with 21 °C during the light hours, and 18 °C during the dark hours. Light intensity was 120 mmol m^-2^s^-1^, and humidity was 70%. Plant transformation was done via the *Agrobacterium tumefaciens* host strain GV3101 *pMP90 pSoup* carrying one of the plasmids in table 1 using the floral dip method (Holsters *et al*., 1980; Koncz and Schell, 1986; Clough and Bent, 1998; Hellens *et al*., 2000; Somssich, 2019). Individual resistant bacteria colonies were selected after two days of growth at 28° C on YT plates (Miller, 1972) supplemented with 50 µg/ml rifampicin, 25 µg/ml gentamycin, 5 µg/ml tetracyclin, and 100 µg/ml spectinomycin, then grown overnight in 250 ml of liquid 2×YT medium with 50 µg/ml rifampicin, 25 µg/ml gentamycin, and 100 µg/ml spectinomycin, shaking at 200 rpm. Cells were harvested by centrifugation at 3200 *g* for 20 min, and resuspended in 300 ml of a 5% sucrose solution with 0.08% Silwet L-77. Plants were then dipped into the solution for approximately 30 seconds with gentle agitation and laid out into a tray covered with cling wrap overnight to maintain humidity. The next day they were returned to upright, and the procedure was repeated after seven days. Once the siliques had ripened, the seeds were harvested and dried for at least two weeks. For the selection of positive transformants, seeds were surface-sterilized using 75% ethanol with 0.1% Triton X-100 on a rotating incubator for at least two hours, after which the seeds were decanted onto filter paper and the ethanol left to evaporate. Once the seeds were dry, they were sprinkled onto a plate of half-strength basal Murashige & Skoog (MS) medium with vitamins and 30 µg/ml hygromycin B, wrapped in aluminum foil, and stratified at 4 °C for three days (Murashige and Skoog, 1962). They were then placed into a growth cabinet for 10 to 14 days, after which healthy looking seedlings were transferred to half-strength MS plates without the hygromycin, and grown for another two weeks, at which stage they were transferred to soil.

### Fungal growth and transformation

The *F. oxysporum* f. sp. *conglutinans* strain 5176 was collected in 1971 from white cabbage (*Brassica oleracea* var. *capitata* (L.)) in Australia (Wang *et al*., 2022*a*). It has since been maintained by the Brisbane Pathology (BRIP) Plant Pathology Herbarium in Queensland, Australia under the accession number BRIP 5176 a. It was first used as a model pathogen for *A. thaliana* by (Campbell *et al*., 2003). To generate a *Fo*5176 line expressing cytoplasmic tdTomato (tdT), an expression clone was generated by cloning the *tdT* coding sequence (amplified from an AddGene-derived plasmid (Shaner *et al*., 2004)) into the *Bgl*II cloning site of plasmid pLAU2 using Gibson assembly (Idnurm *et al*., 2017). This places the gene under the control of a constitutive promoter from the *actin* gene of ascomycete *Leptosphaeria maculans*. After replication in *E. coli*, this plasmid pMAI32 was electroporated into *A. tumefaciens* strain EHA105 with selection on LB medium with kanamycin (50 µg/ml). *Fo*5176 was routinely cultured on potato dextrose agar (PDA) plates. To generate spores for transformation, five plugs of about 5 mm diameter were inoculated into 50 ml half strength potato dextrose broth and cultured at 150 rpm for five days at room temperature, then filtered through miracloth, centrifuged at 3000 g for 5 min, and resuspended in sterile H_2_O. The spores were then transformed with the *A. tumefaciens* strain that had been cultured overnight in LB broth with kanamycin, using standard methods (Idnurm *et al*., 2017). That is, fungal spores and bacteria were mixed on induction medium plates and cocultured for three days. Selection for transformants used an overlay of PDA supplemented with hygromycin and cefotaxime (50 µg/ml and 100 µg/ml, respectively). Transformants that grew through the overlaid medium, were subcultured onto PDA containing hygromycin and cefotaxime and allowed to produce conidia, which were separated with a metal loop to generate strains derived from a single conidium.

### Plant-fungus co-cultivation and infection

To obtain fungal spores, *Fo*5176 was grown for at least seven days on PDA plates at room temperature, at which time five pieces of roughly 1 mm^2^ size were cut from the plate and dropped into a flask containing 50 ml yeast nitrogen base (YNB) medium with 1% sucrose. The liquid cultures were incubated for four days at room temperature with shaking at 120 rpm. The solution was then filtered through miracloth, the spores harvested by centrifugation at 3000 rcf for 10 min and resuspended in 25 ml sterile MilliQ water. For the co-cultivation of plant and fungus, *A. thaliana* seedlings were grown on a vertical petri dish containing half-strength basal MS medium with vitamins for 11 days and then transferred to a horizontal petri dish with a 2-3 cm strip of half-strength basal MS medium with vitamins at the top end, while the rest of the plate was filled roughly 2-3 mm high with liquid quarter-strength basal MS medium with vitamins (this is a setup with slight modifications as described in (Tintor *et al*., 2020)). The seedlings were placed onto the thin MS medium strip at the top end, with the root in the liquid medium. Fungal spores were then added to the liquid medium. The plates were covered with aluminum foil up until the leaves of the plant, and then placed into a growth chamber.

### Microscopy

We imaged the infection and the progression of colonization on a Leica M205 FA stereomicroscope. Infection could usually be observed on day three after spored addition, and at day 5 there was robust colonization. We usually imaged daily from day 3 to 11 dpi. For the fluorescence coming from the plant 2xmT2 reporters, we used the Leica ET CFP (ET436/20x ET480/40m) filter, and for the fungal tdT reporter the Leica ET mCHER (ET560/40x ET630/75m) filter. We use 80× magnification for the images used in this paper. The settings for the imaging (illumination strength, exposure time, gain, etc.) were kept constant for all imaging sessions, to allow for at least semi-quantitative imaging, and to make the images at least relatively comparable. The images were recorded using the Leica Application Suite software, and processed using Fiji Is Just ImageJ (FIJI) and the GNU Image Manipulation Program (GIMP) (Schindelin *et al*., 2012).

## Data availability

We have donated the pGG-PIP vector collection to AddGene (Deposit-ID: 82532, Catalog-#: 196739-196813).

## Acknowledgements

This work was funded by the Australian Research Council (grant no. DE200101560) and was further supported by a seed grant from the Melbourne University Botany Foundation. LW was supported by the China Scholarship Council. H-WC was supported by a Graduate Research Scholarship from the University of Melbourne. Imaging was done on instruments maintained by the Biological Optical Microscopy Platform (BOMP) and the BioSciences Microscopy Unit at the University of Melbourne. The authors would like to thank Dr. Imre E. Somssich for critical reading of the manuscript.

## Supplementary information: Supplementary table 1: List of primers used to clone the promoters

MAS-GG-ACS2p-F AACAGGTCTCAACCTCGGTCGATGTAAATGGATTAAATTTTATA

MAS-GG-ACS2p-R AACAGGTCTCATGTTGCTGTGTCAATTCTCACTTCTTTG

MAS-GG-ACS2pm-F AACAGGTCTCAGGTGTCCTCAAGGTTTCTGTTTCAAC

MAS-GG-ACS2pm-R AACAGGTCTCACACCAAATAAAGTTGATGTGGGGTC

MAS-GG-AOSp-F AACAGGTCTCAACCTACACTTAGACACCCCAATATTTTAGATTT

MAS-GG-AOSp-R AACAGGTCTCATGTTCTATTCGAAACAGTGGCGAGT

MAS-GG-AOSpm1-F AACAGGTCTCAGCCAATATTTTAGATTTTACTTTAAAGAAAT

MAS-GG-AOSpm1-R AACAGGTCTCATGGCGTCTCTAAGTGTTTTTTTTT

MAS-GG-AOSpm2-F AACAGGTCTCAGACACCTAAGTATTTTCTTTCCAACAA

MAS-GG-AOSpm2-R AACAGGTCTCATGTCCTTCTCTTTGAATAAACCTCA

MAS-GG-ARR5p-F AACAGGTCTCAACCTATATGATTTTTTCAAAAGAAAACACCATTTAGT

MAS-GG-ARR5p-R AACAGGTCTCATGTTATCAAGAAGAGTAGGATCGTGACTCGT

MAS-GG-ARR5pm-F AACAGGTCTCAGATTAGGATTATTCTTTATAGAATGTTTTGGTGC

MAS-GG-ARR5pm-R AACAGGTCTCAAATCGTCTCGGTTTTTACCTCTCAAATAGTAT

MAS-GG-ATG8Ap-F AACAGGTCTCAACCTTTAGCAGTGCTTAGTGAGCTTAAATTATAGTT

MAS-GG-ATG8Ap-R AACAGGTCTCATGTTAATTAATAAACTCGATCGTCTGCTAGATCG

MAS-GG-BAK1p-F AACAGGTCTCAACCTGTTTTTTGGAAACAGAGAGAAAACTCA

MAS-GG-BAK1p-R AACAGGTCTCATGTTTATCCTCAAGAGATTAAAAACAAACCCTA

MAS-GG-BIK1p-F AACAGGTCTCAACCTCGTTCCCAAATCTCGGTCAATTG

MAS-GG-BIK1p-R AACAGGTCTCATGTTCAAAGCTAAGAACAGATTCGTTTTCTTCTT

MAS-GG-BIK1pm-F AACAGGTCTCAGTACATTCATTTTGATTGGGATTTATCTTTTTA

MAS-GG-BIK1pm-R AACAGGTCTCAGTACAGACCAAAACAGTGGACTTGGG

MAS-GG-BPS1p-F AACAGGTCTCAACCTAAAAAGAGAGTCACTTGGGTTAAGTGATTT

MAS-GG-BPS1p-R AACAGGTCTCATGTTCTGATAAGTTTCAACTGAAAAAAACAAGAA

MAS-GG-CAD5p-F AACAGGTCTCAACCTGACTTGGGTTCAGTTAAAAATCTCA

MAS-GG-CAD5p-R AACAGGTCTCATGTTTTTGATGATTCTTTCTTCTTTCTTATC

MAS-GG-CERK1p-F AACAGGTCTCAACCTGTATGAAGAAGGGTAACAATTCAACTCTAA

MAS-GG-CERK1p-R AACAGGTCTCATGTTTGAAGCTTCCTTAGATTCCCCAGAGGAAGGGTGTCTGTT

MAS-GG-CIPP1p-F AACAGGTCTCAACCTCAAACCAGACATACTTGCAGCTTCTC

MAS-GG-CIPP1p-R AACAGGTCTCATGTTGAGAGTCAGATGTTCCACAAAAATCTACTC

MAS-GG-CIPP1pm-F AACAGGTCTCAGTTTGCAATCTGTACTAGAATAATATCGTGG

MAS-GG-CIPP1pm-R AACAGGTCTCAAAACGGATCTATATAACATATACGTATGCAATTAA

MAS-GG-CORK1p-F AACAGGTCTCAACCTGTACACTTATTAACATATATTTTTAATTTTGTG

MAS-GG-CORK1p-R AACAGGTCTCATGTTCGTCGACGACCAAAGATGTGAGA

MAS-GG-CORK1pm-F AACAGGTCTCAAGCCTGTTATTCTTAGTATTGCTTTCTATTTAAG

MAS-GG-CORK1pm-R AACAGGTCTCAGGCTTTAAACAATATAAAGCCCACAAG

MAS-GG-CPK29p-F AACAGGTCTCAACCTAATGAGGTAATGGAGGTTATCTTATCGA

MAS-GG-CPK29p-R AACAGGTCTCATGTTGTGAGCAAAGTAGATCGGTCTTCGA

MAS-GG-CPK29pm-F AACAGGTCTCAAGACACCCCCTCCAAGGGGCT

MAS-GG-CPK29pm-R AACAGGTCTCAGTCTCTAGGTTCTCCTATTCCTTTGCAA

MAS-GG-CPK5pro-F AACAGGTCTCAACCTACCATGTGACTACGACAACTACTGG

MAS-GG-CPK5pro-R AACAGGTCTCATGTTGAAACAATGGGAATTACCAAATCC

MAS-GG-CSLD2p-F AACAGGTCTCAACCTGACGACCAAGACTAGAGTTTTGGTTCG

MAS-GG-CSLD2p-R AACAGGTCTCATGTTAGTTAGGATCTAACTTGGCAGATCCCT

MAS-GG-DORN1p-F AACAGGTCTCAACCTGATGTAAAATTTGAAGCTTGAAGATGAAC

MAS-GG-DORN1p-R AACAGGTCTCATGTTGCAGATGATGAATCAGAGAGTCTGG

MAS-GG-EDS16p-F AACAGGTCTCAACCTGACTGCAGAGATCAATTTTCTTTTATTTTAT

MAS-GG-EDS16p-R AACAGGTCTCATGTTGCAGAAATTCGTAAAGTGTTTCTTGA

MAS-GG-EDS16pm1-F AACAGGTCTCACACCGCGTCCAACATTTTAAAACA

MAS-GG-EDS16pm1-R AACAGGTCTCAGGTGTCAGACTCTCAGCTGAACATAATT

MAS-GG-EDS16pm2-F AACAGGTCTCATCTGAAAGAGCCTAAGTGGGTTTCC

MAS-GG-EDS16pm2-R AACAGGTCTCACAGACCAGTTTTTATCATTTAAAAAATATTGTTA

MAS-GG-EDS1p-F AACAGGTCTCAACCTGTTTATCAGATTCCACGTACGATATGTTCTT

MAS-GG-EDS1p-R AACAGGTCTCATGTTGATCTATATCTATTCTCTTTTCTTTAGTGGACTTTC

MAS-GG-EDS1pm1-F AACAGGTCTCACGTGACCAAATCTGAAAACCCAAGT

MAS-GG-EDS1pm1-R AACAGGTCTCACACGTATAGAAGAAATCTACTACTTTAACTCTGTT

MAS-GG-EDS1pm2-F AACAGGTCTCACTCTCATGATGGGGTATTTTGGGTAAC

MAS-GG-EDS1pm2-R AACAGGTCTCAAGAGCATTTCAATGCAAAAATGGGT

MAS-GG-EFRp-F AACAGGTCTCAACCTATCTAGACGATTAAGTAATTGAGCATGTAAAAG

MAS-GG-EFRp-R AACAGGTCTCATGTTGTCGATTATAAAAAGATAAAAGAAAGGTTCTTCT

MAS-GG-ELI3p-F AACAGGTCTCAACCTTAAAGTCGATGTTCTATATGTATTCAAAATAAT

MAS-GG-ELI3p-R AACAGGTCTCATGTTATGGATAAATAATAAGCGAATGGGA

MAS-GG-ERF1p-F AACAGGTCTCAACCTCTCTCCCAATTGATATTTTTGTTATTTCT

MAS-GG-ERF1p-R AACAGGTCTCATGTTGTAGAAAAAATACTCTGTTTCTTGACTACTCTGT

MAS-GG-EX1p-F AACAGGTCTCAACCTCCGCCGTCTTAAGTGGAATTTGG

MAS-GG-EX1p-R AACAGGTCTCATGTTCGCCGGAGAGATGTGAGAGCG

MAS-GG-FERp-F AACAGGTCTCAACCTAGAAAAGTTAAGAGTGGGAACTGGGA

MAS-GG-FERp-R AACAGGTCTCATGTTCGATCAAGAGCACTTCTCCGG

MAS-GG-FH16p-F AACAGGTCTCAACCTCTGACCATAGTGTGGTAACTCAGATATTTT

MAS-GG-FH16p-R AACAGGTCTCATGTTGGATCAGGAACCAAACAGATGATT

MAS-GG-FLS2p-F AACAGGTCTCAACCTTTTTTGTTAGAGAGATTTTTGTTGTTTTTGTT

MAS-GG-FLS2p-R AACAGGTCTCATGTTGGTTTAGACTTTAGAAGAGTTGAAATTGTGG

MAS-GG-FMO1p-F AACAGGTCTCAACCTCGGAAAAATCCTTCGTCAATGTGTG

MAS-GG-FMO1p-R AACAGGTCTCATGTTCTGAGAGAGGTTATGCTAGAGAGAAGAGAG

MAS-GG-FRK1p-F AACAGGTCTCAACCTAAATTAAACGCCTTTTTATCAACAAC

MAS-GG-FRK1p-R AACAGGTCTCATGTTACTTAATTGAGCTGCTTTCTCTGG

MAS-GG-FRK1pm-F AACAGGTCTCACGTCTCTTTACATTTGTGATGTGGT

MAS-GG-FRK1pm-R AACAGGTCTCAGACGAACACTGATATAAAAAATCTCACA

MAS-GG-GLR25p-F AACAGGTCTCAACCTGATAAGAAGTGATTCAGCTGGGGTTT

MAS-GG-GLR25p-R AACAGGTCTCATGTTATTGATAGCTCGCAAGCTCAAATCTG

MAS-GG-GLR25pm-F AACAGGTCTCAACCTCTCTCACCCTGGTCGCAAA

MAS-GG-GLR25pm-R AACAGGTCTCAAGGTTTTCTCTTTTCAACGTACAATATTTACATGA

MAS-GG-GLR27p-F AACAGGTCTCAACCTCACACACTGGTCACTTAATGGTTTATTTGA

MAS-GG-GLR27p-R AACAGGTCTCATGTTCCAGATTGAGGAACTTTATGTCATTCTTTAAC

MAS-GG-GLR27pm-F AACAGGTCTCAACTCAAAATGATGCGTATCATTAATTCTATAGTTT

MAS-GG-GLR27pm-R AACAGGTCTCAGAGTTTGAAATTTTTAACAAAAGTCTTAAGTTATATCAG

MAS-GG-GPAT5p-F AACAGGTCTCAACCTAAAAAGCGTTTTAATTAGAGAGATTTTTGC

MAS-GG-GPAT5p-R AACAGGTCTCATGTTCTTTTGTTTTTTGCTCGAATATTATTTT

MAS-GG-HPCA1p-F AACAGGTCTCAACCTAAACATAAGAGAAAACGCAAGTTGATGA

MAS-GG-HPCA1p-R AACAGGTCTCATGTTCTTCAAACCCAAAAAGAACCTCTTATCA

MAS-GG-HRMp-F AACAGGTCTCAACCTCTGAATATCTTCTTTTGGTTGCTCTGA

MAS-GG-HRMp-R AACAGGTCTCATGTTCGTAATATCTCTCTGTTTTTGCTCTGTTTT

MAS-GG-LECRKVI2p-F AACAGGTCTCAACCTACAAAGTCAATTACTTCGAGTTTTTTTCTG

MAS-GG-LECRKVI2p-R AACAGGTCTCATGTTGGGTGAGCGAAGTAAAGAAGGAGATA

MAS-GG-LYK5p-F AACAGGTCTCAACCTATTTTCTGTTAAGTTTGAACATTTGGTTGTAA

MAS-GG-LYK5p-R AACAGGTCTCATGTTTTGTGGTGTTCTGATCTGAAGAGG

MAS-GG-LYK5pm-F AACAGGTCTCACAACAGACCAAGACCATCTTTATGTCC

MAS-GG-LYK5pm-R AACAGGTCTCAGTTGGATCTCATGTGAAAGAGACACATT

MAS-GG-LYM1p-F AACAGGTCTCAACCTATATCATCGGTAAGTCACTAGACTATTGAACG

MAS-GG-LYM1p-R AACAGGTCTCATGTTTTGTGTTTAGGGTTTTACGAAATTCAA

MAS-GG-MIK2p-F AACAGGTCTCAACCTGTAAATAACGTTGAACTCGCGG

MAS-GG-MIK2p-R AACAGGTCTCATGTTACAGTTGCAGATTATCTCTCTACGGTC

MAS-GG-MLO6p-F AACAGGTCTCAACCTAAATATACATTTGGTTGACATGTTTCTCATT

MAS-GG-MLO6p-R AACAGGTCTCATGTTAGAACTCACAGAACAGTTCCAAGCAAA

MAS-GG-MPK3p-F AACAGGTCTCAACCTAAAAAAATTCTGATCGAAAATAGCTTAC

MAS-GG-MPK3p-R AACAGGTCTCATGTTCTCTCTCAATTGATCAAAGTCGA

MAS-GG-MPK4p-F AACAGGTCTCAACCTGACTTGTTTGTGAATATAGAGGAAACATGTAATTAT

MAS-GG-MPK4p-R AACAGGTCTCATGTTCGGAGCAAAATTCCTCACAACAACG

MAS-GG-MPK6p-F AACAGGTCTCAACCTAACACAAGAGAAGAGATTTATTGCTTC

MAS-GG-MPK6p-R AACAGGTCTCATGTTGACCGGTAAAGATGAAAGCTTTT

MAS-GG-MYB15p-F AACAGGTCTCAACCTGATGAATTTGAATAAACTAAACAAAATT

MAS-GG-MYB15p-R AACAGGTCTCATGTTCTCTTTGATTTGTGATTGCTGATAAA

MAS-MYB15m-F AGAGGACCATGGACACCTGAAGAAGATCAAATCTTT

MAS-GG-MYB72p-F AACAGGTCTCAACCTACACGATCTCTTTTGAGATTTAAGAAG

MAS-GG-MYB72p-R AACAGGTCTCATGTTCTTATTACACTACTTTCTTCTCTATAGCTACC

MAS-GG-MYB72pm1-F AACAGGTCTCACGTTTTAAAACTTTACCTTATGTCCAATCTCT

MAS-GG-MYB72pm1-R AACAGGTCTCAAACGTGACGTAGCATGTGTGGGTC

MAS-GG-MYB72pm2-F AACAGGTCTCAGCTCTCTCTCTACGAGTGAAGTGCCT

MAS-GG-MYB72pm2-R AACAGGTCTCAGAGCCAAAAGCATGGAACGTACG

MAS-GG-NET4Ap-F AACAGGTCTCAACCTTTAATCCTCTTCTCGTACATCACAT

MAS-GG-NET4Ap-R AACAGGTCTCATGTTGGCTGCAAAAATCAATGGACC

MAS-GG-PAD4p-F AACAGGTCTCAACCTAATTAGGGTTTTATCAGATTAAAGAGATTTACTGATT

MAS-GG-PAD4p-R AACAGGTCTCATGTTGATTGGATATCGAGTAGAGAGTTGCAGA

MAS-GG-PDF12p-F AACAGGTCTCAACCTTCTACCAAAAATCTTTGGTGCTTGATC

MAS-GG-PDF12p-R AACAGGTCTCATGTTGATGATTATTACTATTTTGTTTTCAATGTATAGA

MAS-GG-PEP1p-F AACAGGTCTCAACCTGAAGTCAAAAATTGAGTCGAAAAATC

MAS-GG-PEP1p-R AACAGGTCTCATGTTGAGATCTGATAAGACAGAGGAAAACTT

MAS-GG-PEP2p-F AACAGGTCTCAACCTTGAAGCTCTTGTGAATAGAGAAGAGA

MAS-GG-PEP2p-R AACAGGTCTCATGTTGAAATCCAATAGTTTGGTGAGTTATC

MAS-GG-PEP3p-F AACAGGTCTCAACCTGCACTTTAAGTTACATTGTTTAGTCTAATTATT

MAS-GG-PEP3p-R AACAGGTCTCATGTTCGTTGACTTCTTAATCTTTTTTTGGGAA

MAS-GG-PEPR1p-F AACAGGTCTCAACCTAGAGAAGGAAAACAACCATGTATTCCAG

MAS-GG-PEPR1p-R AACAGGTCTCATGTTCTGAGTTTAAAGATCGAGAAACATGCAG

MAS-GG-PEPR1pm-F AACAGGTCTCAGAAACCAAACATCTCGTCATAAAAAAC

MAS-GG-PEPR1pm-R AACAGGTCTCATTTCTCTGTATACCAACGATTGTGAGA

MAS-GG-PEPR2p-F AACAGGTCTCAACCTAGTTTGAGATGGAGTTGCATTGTG

MAS-GG-PEPR2p-R AACAGGTCTCATGTTGAGATTAGAGCTCAAGAGACTGAAATAT

MAS-GG-PER5p-F AACAGGTCTCAACCTCAGTGCGTAGTAGTGAGTTTTCTTCA

MAS-GG-PER5p-R AACAGGTCTCATGTTATTTGTAGATCTCACTTGGTATATATTTCGTAC

MAS-GG-PER5pm-F AACAGGTCTCAGACGAATATATATAATTAGCTACTAAATTAAATT

MAS-GG-PER5pm-R AACAGGTCTCACGTCTCAGAACGAGTGAATGATTC

MAS-GG-PLP1p-F AACAGGTCTCAACCTCTGATCATCTAGCCTCTTCCC

MAS-GG-PLP1p-R AACAGGTCTCATGTTAATAGTTGATCGATCTTCTTTTGAGTTAA

MAS-GG-PMR4p-F AACAGGTCTCAACCTGCTCGATGTCGATTTGAGACGTAGT

MAS-GG-PMR4p-R AACAGGTCTCATGTTAGTAGCATGTGGTAGATCTTAGAAATTTCTCG

MAS-GG-PR1pm-F AACAGGTCTCACTCCCTCCATATAAAAAAGTTTGATTTTATAG

MAS-GG-PR1pm-R AACAGGTCTCAGGAGAATCATTTTATAAGTTAAAACAAGCTTG

MAS-GG-PR1pro-F AACAGGTCTCAACCTATATATAACGATCATTGATTAGTATATATACATATTG

MAS-GG-PR1pro-R AACAGGTCTCATGTTTTCTAAGTTGATAATGGTTATTGTTGT

MAS-GG-RALF23p-F AACAGGTCTCAACCTGGTGATTCCGGTTTCCGACG

MAS-GG-RALF23p-R AACAGGTCTCATGTTTCTTCTGTACACTGTAGCTTTAGCTCTCTC

MAS-GG-RALF23pm-F AACAGGTCTCAGAGCACTCATAATTGTACAAAATAAAAGTAAATG

MAS-GG-RALF23pm-R AACAGGTCTCAGCTCTCCTTCCATGATTTGAGACTATTTC

MAS-GG-RBOHDp-F AACAGGTCTCAACCTGACTTGTTAAATTGCTCTCTTAGTCTTA

MAS-GG-RBOHDp-R AACAGGTCTCATGTTCGAATTCGAGAAACCAAAAAGATC

MAS-GG-RBHOFp-F AACAGGTCTCAACCTACCGGTTGAAAATAAGAGTGGTGGA

MAS-GG-RBOHFp-R AACAGGTCTCATGTTAGATCCAAAGTCGGAATTCAAAGAGTT

MAS-GG-RBOHFpm-F AACAGGTCTCATGCAGAAGATAGTGAAGATAGTTGCAGAA

MAS-GG-RBOHFpm-R AACAGGTCTCATGCAACTTTTATAGTTTTCGAACGAAAGTA

MAS-GG-RCD1p-F AACAGGTCTCAACCTGGAGGAGCAGATTGGACACCGT

MAS-GG-RCD1p-R AACAGGTCTCATGTTCTATATATTAACAATACTAAACCTATAACCTTGATAG

MAS-GG-RCD1pm-F AACAGGTCTCAATACGTCTCATATAGTTATGCTGATTCTTTCTTG

MAS-GG-RCD1pm-R AACAGGTCTCAGTATGATCCTGTAATATCATTCCTTCACAAAA

MAS-GG-RFO1p-F AACAGGTCTCAACCTATATTAACCATGCATGCAAACAAA

MAS-GG-RFO1p-R AACAGGTCTCATGTTTTTTTTTCTCTAATGACTTTTATGTATG

MAS-GG-RLP26p-F AACAGGTCTCAACCTGATTAAAGGATTGATCGGTAAACAAC

MAS-GG-RLP26p-R AACAGGTCTCATGTTGGTGTTTGTGATTGAACCAACAAGT

MAS-GG-RLP29p-F AACAGGTCTCAACCTCCAGCAAAAAGCTTCTTCTACTCAA

MAS-GG-RLP29p-R AACAGGTCTCATGTTAGGTTTTGGTGTAAGAGAGAGGAAAGA

MAS-GG-RPS4p-F AACAGGTCTCAACCTCGAGAACCTTGGCGAACTTGTCA

MAS-GG-RPS4p-R AACAGGTCTCATGTTGGCCCAAAAGCTTTTTCCCGGT

MAS-GG-SCOOP12p-F AACAGGTCTCAACCTAATAGGTTTCGAGTACTGTATTGATGTTTAACTG

MAS-GG-SCOOP12p-R AACAGGTCTCATGTTCTCGATCTTTATTTTTTTCTCGAGTTTAGA

MAS-GG-SOBIR1p-F AACAGGTCTCAACCTTTTCGATTTTTCTAATCTCACAGCTGTTC

MAS-GG-SOBIR1p-R AACAGGTCTCATGTTTAATTAGAGAAAGTTTCTTCTTGTGGATGTT

MAS-GG-SULTR41p-F AACAGGTCTCAACCTATGATCCATCACACGCCTGCCT

MAS-GG-SULTR41p-R AACAGGTCTCATGTTGATGGCTCTTGCGCACGCTTGG

MAS-GG-SULTR42p-F AACAGGTCTCAACCTGTAGCTTCCACGCCCTTGCCTAA

MAS-GG-SULTR42p-R AACAGGTCTCATGTTCGGAATTGGTGGGATAGAGAAGAAT

MAS-GG-TET8p-F AACAGGTCTCAACCTCGGATGTATCAAAGGTAAAAATATC

MAS-GG-TET8p-R AACAGGTCTCATGTTGGTTTAGATTCAGAGAGAAAGATTG

MAS-GG-TUB6p-F AACAGGTCTCAACCTATTTAGAGGGTGTTATTGGTTTGTG

MAS-GG-TUB6p-R AACAGGTCTCATGTTCTTCTATTTTATCTGAAATCAACATTACA

MAS-GG-TUB6pm1-F AACAGGTCTCATAACAAAAAGTTATGAATATTCACAGACATA

MAS-GG-TUB6pm1-R AACAGGTCTCAGTTATGGTTAACCGAGGATGAGC

MAS-GG-TUB6pm2-F AACAGGTCTCAGAGGCCATTTTTTTTTCCCGT

MAS-GG-TUB6pm2-R AACAGGTCTCACCTCATTGCGTATGACAATGCG

MAS-GG-VSP2p-F AACAGGTCTCAACCTTCTCTCTGGTTATATTTTGTTGCTGCTT

MAS-GG-VSP2p-R AACAGGTCTCATGTTGTTTTTTATGGTATGGTTTATTGTTTAGTTTGTG

MAS-GG-WAKL22p-F AACAGGTCTCAACCTATATTAACCATGCATGCAAACAAA

MAS-GG-WAKL22p-R AACAGGTCTCATGTTTTTTTTTCTCTAATGACTTTTATGTATG

MAS-GG-WRKY11p-F AACAGGTCTCAACCTTAGTTCCAAAACCGCATTGACAT

MAS-GG-WRKY11p-R AACAGGTCTCATGTTGATGATTTCTTGGTCTGAGGATTTT

MAS-GG-WRKY11pm-F AACAGGTCTCAGACGAAACTGTTGATTGCTTTATTCC

MAS-GG-WRKY11pm-R AACAGGTCTCACGTCTCCTCAAAGTTCGAGGTTACT

MAS-GG-WRKY17p-F AACAGGTCTCAACCTGTCTCGCAGAGGTTATTTATCTACTTGGTT

MAS-GG-WRKY17p-R AACAGGTCTCATGTTGATGAGAAACCAGAGGAGAAACTTGAAG

MAS-GG-WRKY17pm1-F AACAGGTCTCAGGTCAACGATTCCCATGTCGCTAA

MAS-GG-WRKY17pm1-R AACAGGTCTCAGACCTAACCGACTAATATATATGATTGTCGTG

MAS-GG-WRKY17pm2-F AACAGGTCTCAAAGCAGACCAAACTTTGATTACTTTATTCCATA

MAS-GG-WRKY17pm2-R AACAGGTCTCAGCTTGAGTTGTGAGATATGTAGGGTCTTCTT

MAS-GG-WRKY33p-F AACAGGTCTCAACCTCGCTGCTTTTTCGAGATAGATAG

MAS-GG-WRKY33p-R AACAGGTCTCATGTTACGAAAAATGGAAGTTTGTTTTATAA

MAS-GG-WRKY40p-F AACAGGTCTCAACCTTGTGTATAACTATTATGCAGCCTTTTTCAA

MAS-GG-WRKY40p-R AACAGGTCTCATGTTGTAAATATATGTAGGATGAATCTTCGATATGGGT

MAS-GG-WRKY40pm-F AACAGGTCTCATACAAGAATAGGTACAGTCCTGGTTTGTG

MAS-GG-WRKY40pm-R AACAGGTCTCATGTAATTGTGAATAATAAAATCTTAATTCAGAT

MAS-GG-WRKY53p-F AACAGGTCTCAACCTATCTTGTGAGCTGATTCAAAGATTTC

MAS-GG-WRKY53p-R AACAGGTCTCATGTTTTAGTATATGATTCCCAAAATAGATTTTTT

MAS-GG-WRKY70p-F AACAGGTCTCAACCTCATTGTAGATATGATATATGAAGCTTCCCC

MAS-GG-WRKY70p-R AACAGGTCTCATGTTGTTAGTTTTGAGGAAGTTTTTGGTGAG

MAS-GG-WRKY70pm-F AACAGGTCTCAGTATCTCGCATATTAACTTAGGCTAGAGAGC

MAS-GG-WRKY70pm-R AACAGGTCTCAATACTATGATAAACCAGTTGGTTCTGTAGCG

MAS-GG-XLG2p-F AACAGGTCTCAACCTGAGTGGAGGAGCATAGTGTGATTATTTAC

MAS-GG-XLG2p-R AACAGGTCTCATGTTCTTCTTACCCAATCAAGCACACATACAA

**Figure S1:**
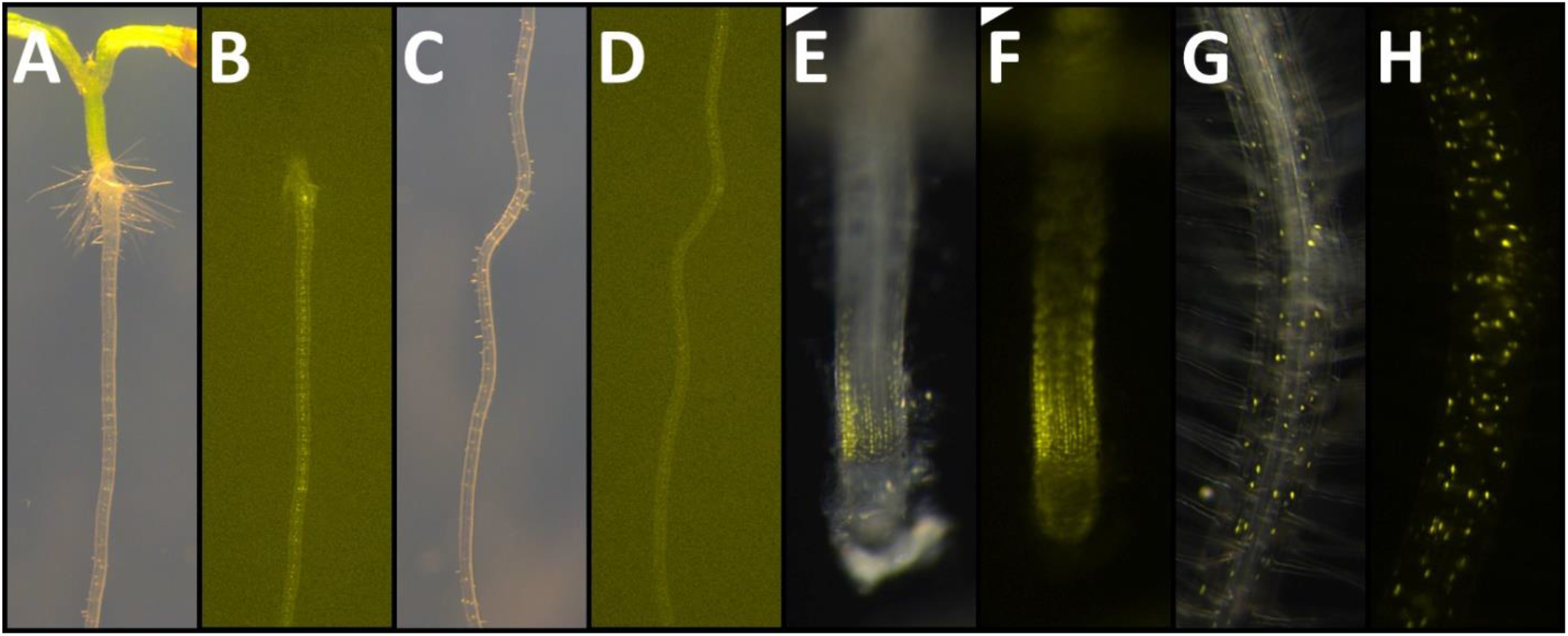
Expression pattern of *PEP1* and *PEP2*. **A-D** shows the expression of *PEP1* (yellow). Robust expression is found in the inner root tissues of the mature DZ (A, B). From the young to mature DZ, expression gradually becomes stronger (C, D). No expression in the root tip, MZ or EZ. **E-H** shows the expression of *PEP2* (yellow). Expression is found in the EZ of the tip, then disappears in the young DZ (E, F), and returns to all tissues in the mature DZ (G, H). A, C, E, G are bright field images plus fluorescence, B, D, F, H are fluorescence only.

**Figure S2:**
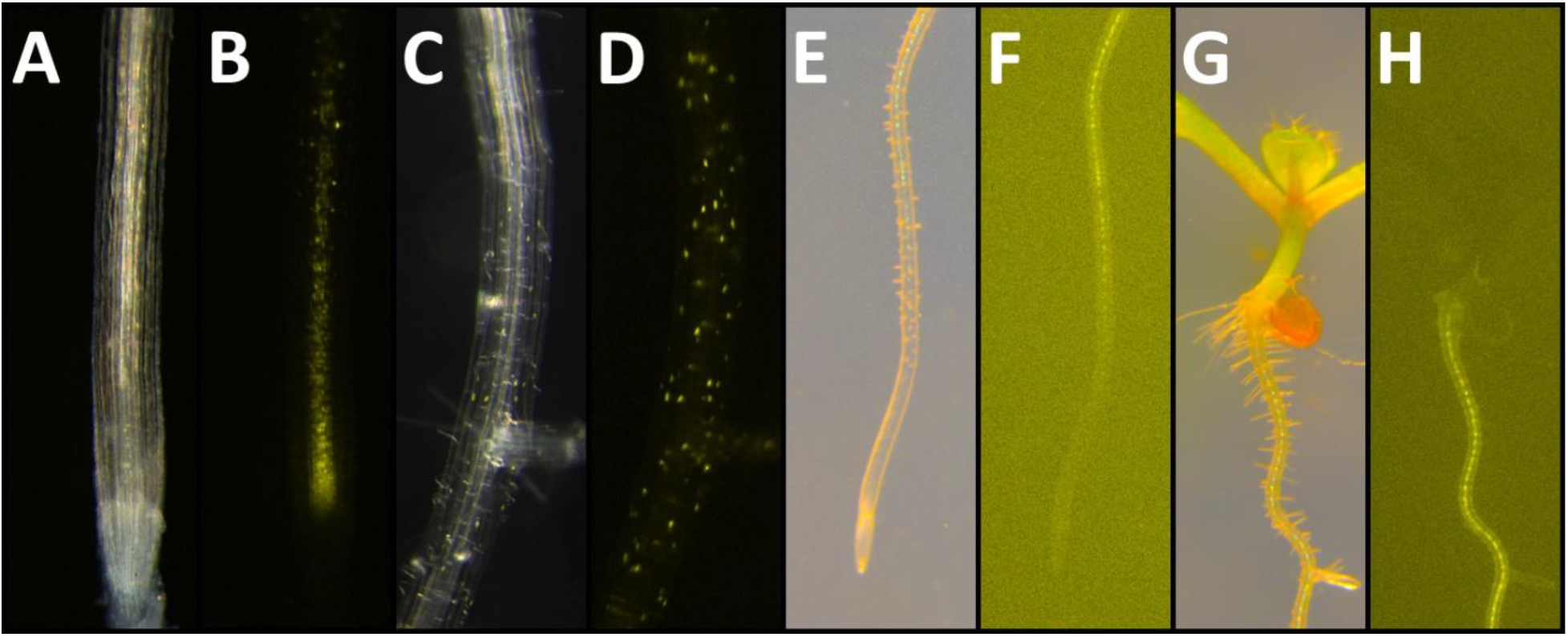
Expression pattern of *PEPR1* and *PEPR2*. **A-D** shows the expression of *PEPR1* (yellow). Weak expression is found in vasculature of the MZ, EZ and young DZ (A, B). In the mature DZ, weak expression is found in all tissues (C, D). **E-H** shows the expression of *PEPR2* (yellow). Strong expression is found in the vasculature of the DZ, starting in the root hair zone (E, F) and becoming stronger in the mature DZ (G, H). A, C, E, G are bright field plus fluorescence, B, D, F, H are fluorescence only.

**Figure S3:**
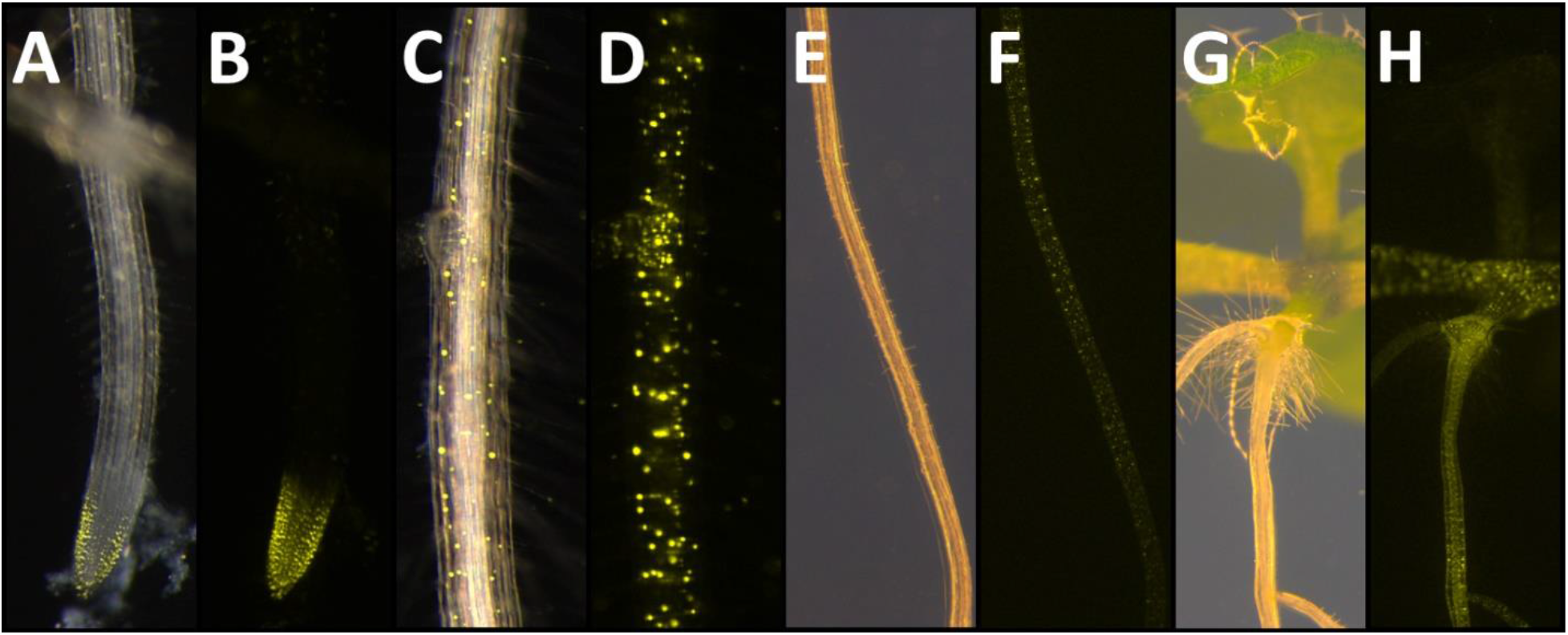
Expression pattern of *BIK1* and *RBOHD*. **A-D** shows the expression of *BIK1* (yellow). Expression is found in the root tip around the meristem, including the root cap and part of the EZ (A, B). Expression appears stronger in the outer tissues compared to the vasculature. Further up the root, expression is robust in the mature DZ, but still stronger in the outer tissues (C, D). **E-H** shows the expression of *RBOHD* (yellow). *RBOHD* is expressed in all cells and tissues from the young DZ onwards (E, F). Expression is strongest in differentiated tissue (G, H), A, C, E, G are bright field plus fluorescence, B, D, F, H are fluorescence only.

